# Evaluation of nanopore sequencing for epigenetic epidemiology: a comparison with DNA methylation microarrays

**DOI:** 10.1101/2022.03.01.482537

**Authors:** Robert Flynn, Sam Washer, Aaron R Jeffries, Alexandria Andrayas, Gemma Shireby, Meena Kumari, Leonard C Schalkwyk, Jonathan Mill, Eilis Hannon

**Author notes:** Correspondence to: Eilis Hannon, University of Exeter Medical School, RILD Building, Royal Devon and Exeter Hospital, Barrack Road, Exeter. EX2 5DW. UK.: NA.

## Abstract

Most epigenetic epidemiology to date has utilized microarrays to identify positions in the genome where variation in DNA methylation is associated with environmental exposures or disease. However, these profile less than 3% of DNA methylation sites in the human genome, potentially missing affected loci and preventing the discovery of disrupted biological pathways. Third generation sequencing technologies, including Nanopore sequencing, have the potential to revolutionise the generation of epigenetic data, not only by providing genuine genome-wide coverage but profiling epigenetic modifications direct from native DNA. Here we assess the viability of using Nanopore sequencing for epidemiology by performing a comparison with DNA methylation quantified using the most comprehensive microarray available, the Illumina EPIC array. We implemented a CRISPR-Cas9 targeted sequencing approach in concert with Nanopore sequencing to profile DNA methylation in three genomic regions to attempt to rediscover genomic positions that existing technologies have shown are differentially methylated in tobacco smokers. Using Nanopore sequencing reads, DNA methylation was quantified at 1,779 CpGs across three regions, providing a finer resolution of DNA methylation patterns compared to the EPIC array. The correlation of estimated levels of DNA methylation between platforms was high. Furthermore, we identified 12 CpGs where hypomethylation was significantly associated with smoking status, including 10 within the AHRR gene. In summary, Nanopore sequencing is a valid option for identifying genomic loci where large differences in DNAm are associated with a phenotype and has the potential to advance our understanding of the role differential methylation plays in the aetiology of complex disease.

## Introduction

There is increasing interest in the role of epigenetic variation in health and disease, with the primary focus of epigenetic epidemiology being on variable DNA methylation (DNAm)(1). The development of standardized assays (e.g. the Illumina Infinium Methylation EPIC BeadChip (“EPIC array”)) have enabled epigenome-wide association studies (EWAS) to identify specific positions in the genome where methylomic variation is associated with environmental exposures or disease. The most common approach for profiling methylomic variation involves the sodium bisulfite treatment of DNA, to differentiate methylated cytosines, which are protected and remain as a cytosine, from unmethylated cytosines, which are converted to uracils. The methylation status at individual genomic positions is then determined by either sequencing the bisulfite-converted DNA or hybridising to a microarray. DNAm level is estimated at individual genomic positions as the proportion of methylated cytosines, which represents the proportion of cells in the sample that are methylated at that position. One of the limitations with using a microarray is that the specific sites profiled is predefined, and in the case of a commercial product such as the EPIC array, predominantly non-customisable. Despite the EPIC array being the most extensive array available, it only captures *3% of CpGs across the human genome (2) and while it assays >97% of RefSeq genes there is a huge range in the number of sites overlapping each gene, with a median of 18 sites per gene. It is highly probable, therefore, that many of the specific sites at which aberrant DNAm underpins the development of a given disease are either not included or weakly indexed by proximal sites in existing analyses. Alternatively, a sequencing based approach will provide a more comprehensive view of the methylome, and is applicable for the study of any organism, with whole genome bisulfite sequencing currently regarded as the gold standard experimental approach (3). A consequence of the bisulfite conversion step is the requirement for bespoke alignment tools as cytosines in the reference genome could generate either a cytosine in sequencing data, representing an methylated site, or a thymine, representing an unmethylated site; this means that a relatively large number of reads have to be discarded at this processing stage due to the inability to assign them unambiguously to the reference genome(4). With any sequencing technology, the accuracy of the quantification of DNAm is dependent on the number of reads overlapping a given genomic position. Given the often stochastic nature of sequencing coverage and the fact that it is effectively count data, having sufficient depth at any one site in all or even the majority of your samples is often unlikely making it unfeasible with the current technologies to perform EWAS in large population cohorts.

Sequencing technologies continue to evolve, with novel long-read approaches being able to interrogate epigenetic modifications, including DNAm, in parallel to determining the underlying DNA sequence. This bypasses the need to perform a bisulfite treatment on the DNA. For example, Oxford Nanopore Technologies (ONT) sequencing platforms use known electrical signal profiles to call nucleotide bases from DNA fragments, which can be further refined to distinguish methylated cytosine from unmethylated cytosine(5). While the application of these technologies to large populations is primarily limited by their cost, it has yet to be established whether the quantification of DNAm is sufficiently accurate to detect differentially methylated sites in an epidemiological study, and how the estimation compares to the standard microarray technology.

In order to increase the likelihood of obtaining sufficient coverage in the same regions of the genome, a targeted approach coupled with sequencing can be used. There are a number of existing approaches for targeting specific regions in bisulfite-based sequencing, but Nanopore sequencing can detect DNAm directly from sequence data that has not been bisulfite-converted. Nanopore Cas9-targeted sequencing (nCATS) is one such method which uses Cas9/guide RNA (gRNA) ribonucleoprotein complexes (RNP) to selectively cut DNA around the targeted region and enrich these regions prior to sequencing (6). While this has been shown to be effective at increasing the depth of sequencing in these regions, it is still unclear whether this will confer sufficient sensitivity when quantifying the level of DNAm such that differences between groups can be detected. This is vital for assessing whether this could be a plausible approach for epigenetic epidemiology studies of complex traits, which are typically associated with small differences in DNAm between groups.

One of the phenotypes with the most dramatic influence on DNAm profiles is tobacco smoking, where the signature is not only detectable in the blood of current and former smokers (7–9), but additionally in the blood of new-borns and children who were exposed in utero (10–12). In the largest meta-analysis comparing 2,433 current and 6,956 never smokers, 2,623 DNAm sites, annotated to 1,405 genes were identified with significantly different levels of DNAm, many of which were associated with large effects (> 5%) (13). Harnessing DNAm levels at multiple sites into an aggregate score has been shown to be highly predictive of current smoking status (7, 14).

The aim of this study was to assess the viability of using targeted ONT sequencing for epigenetic epidemiology by attempting to rediscover known differentially methylated positions (DMPs) that existing technologies have shown to be robustly associated with tobacco smoking. We selected three genomic regions containing highly significant smoking-associated DMPs (AHRR, GFI1 and an intergenic region on chromosome 2) and implemented the nCATS methodology, a CRISPR-Cas9 targeted sequencing approach. We report the first comparison of DNAm called from ONT long read data with DNAm profiled using the EPIC array on the same samples and the first assessment of the sensitivity of DNAm quantification with ONT to detect tobacco smoking associated differentially methylated positions by comparing estimated levels of DNAm between a smoker and non-smoker.

## Results

We targeted three genomic regions where previous studies have identified multiple differentially methylated sites associated with tobacco smoking; two are centred on specific genes (AHRR and GFI1) and one was intergenic on chromosome 2q37.1 (**Table 1**). To enrich for reads in these regions we designed a panel of 18 gRNAs (**Supplementary Table 1**) targeting the start and ends, with additional gRNAs tiled across the larger AHRR region (*140kb) optimising the spatial distribution (mean distance between guides = 15.6kb) against the predicted performance based on sequence content. After sequencing two MinION r9.4.1 flowcells on a Nanopore Mk1b sequencer, 215,829 reads were generated. These were aligned to the human genome (hg38) using minimap2 and filtered resulting in 185,540 (86%) high quality primary alignments (**Supplementary Table 2**). Of these, 645 reads (0.35%) were located within our three targeted regions, meaning that all regions had elevated coverage (range of means across regions 7.72 - 21.5) compared to the mean read depth genome-wide (1.18, SD = 0.99), as desired. On closer inspection, acute increases in read depth were observed at the location of all gRNAs, with accumulative effects observed where multiple gRNAs are located within the range of the typical read sizes (**Figure 1**, **Supplementary Figures 1-2**). While it was evident that all gRNAs had successfully targeted the desired genomic locations, the performance in terms of number of reads at each position was variable, in line with random mixing of the gRNAs within the pools (see **Methods**). The proportion of on-target reads was in line with previous studies(6) and off-target reads were randomly distributed across the genome (**Supplementary Figure 3**). The read lengths within the two smaller regions were determined by the size of the region and the location of the gRNA, for example, within the chromosome 2 region, 40.4% of the reads spanned at least 90% of the targeted region (**Supplementary Figure 4**). In contrast, while the larger AHRR region was associated with longer reads, (mean = 10,067bp; **Supplementary Figure 5**) with 160 (38%) of reads longer than 10kb, the proportion of the targeted region captured by a single read was smaller on average (mean = 0.07).

**Table 1.** Summary of the three targeted regions.

| Gene | Chr | Range (hg38) | Size(kp) | Number of guides | Number of reads | % of reads | Mean coverage | Mean read length | N reads > 10kb |
| --- | --- | --- | --- | --- | --- | --- | --- | --- | --- |
| AHRR | 5 | 300325 - 440842 | 140.5 | 10 | 421 | 0.23% | 19.1 | 10067 | 160 |
| - | 2 | 233280458 - 233288795 | 8.3 | 4 | 89 | 0.05% | 21.5 | 4987 | 3 |
| GFI1 | 1 | 92939875 - 92957522 | 17.6 | 4 | 135 | 0.07% | 7.72 | 3380 | 14 |

**Abbreviations**

**Figure 1.**
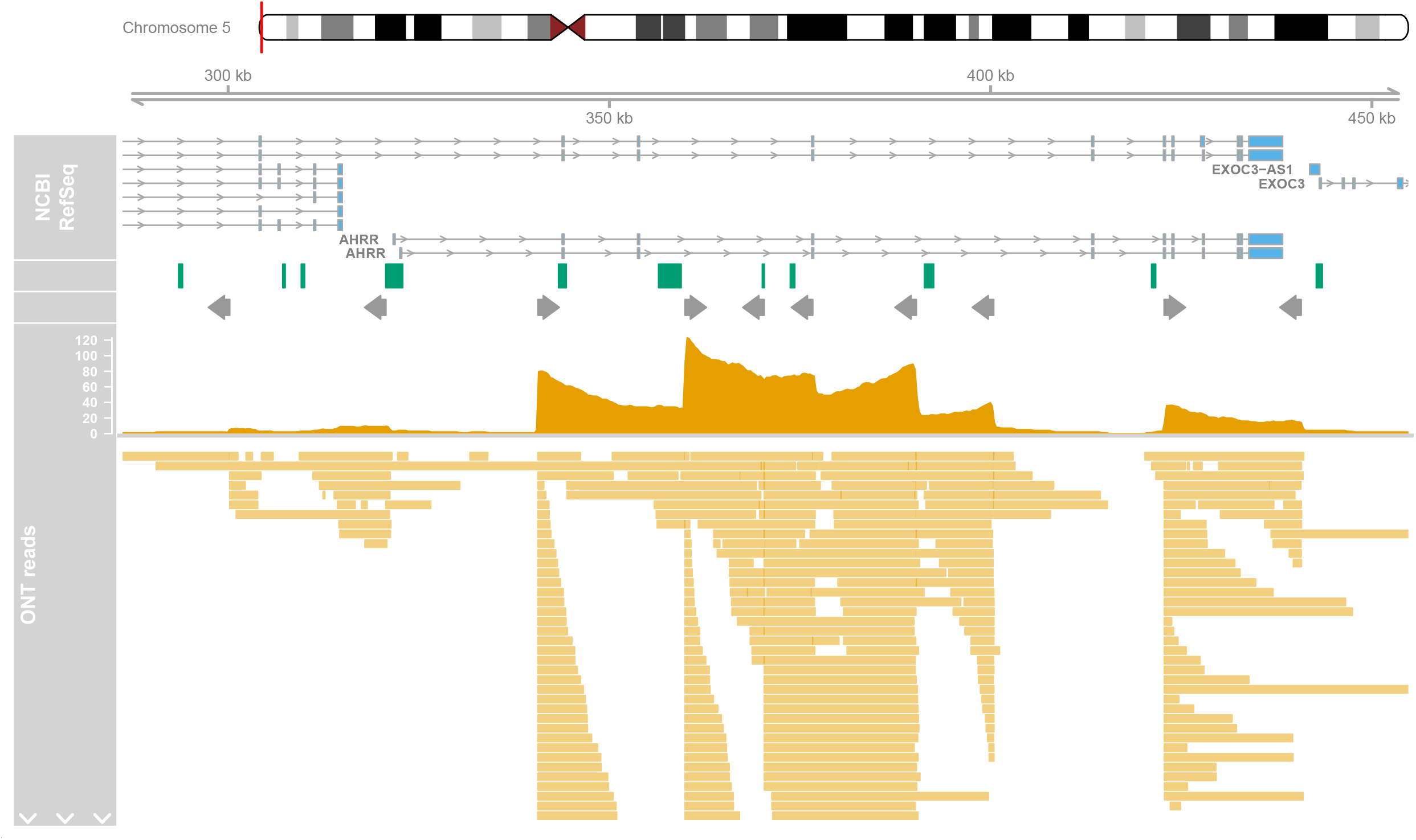
Distribution of nanopore sequence reads across AHRR region. Depicted is the targeted genomic region on chromosome 5 containing AHRR and the location of the gRNAs and sequencing reads. Shown from top to bottom is the gene locations (exons and introns) for different transcripts, CpG islands locations (green boxes), target position of the gRNAs where the grey arrow indicates the orientation, a histogram of the total number of Nanopore reads overlapping each position, and the location of the individual reads at the bottom. Note that due to the high number of reads in the region, only a subset are included to give a representative view of read mappings.

To quantify the level of DNAm across the targeted regions, Nanopolish(5), which uses a Hidden Markov model and the electrical signal data to determine the methylation status at CpG sites was run. Filtering to sites with a minimum read depth of at least 10, DNAm was quantified at 1,779 CpGs clustered into 1,130 regions. This represents a much finer resolution of data than is obtained using the most comprehensive microarray available. For example in the AHRR region, we captured 1,429 CpGs compared to 159 DNAm sites included on the EPIC array, representing *9 fold increase of data points. Furthermore, the median spacing between CpGs in this region is reduced to 35bp in the ONT data compared to 405bp on the EPIC array. First, we were interested in assessing the level of accuracy in the quantification of DNAm from ONT sequencing, by comparing the level of DNAm at sites profiled using the EPIC array. A total of 98 CpGs within the three targeted regions across the two samples were quantified with both platforms. Estimated DNAm levels correlated strongly (r = 0.94; **Figure 2**), although the absolute difference between the two technologies was moderate (RMSE = 0.138). Of note, it appears that the ONT-derived levels of DNAm are less similar between platforms at the extremes; rather than reflecting inaccuracies in the ONT approach we hypothesize that this reflects the fact that in these parts of the distribution the EPIC array is known to be less sensitive(15) with variation here being attributable to lack the of precision in the array derived estimates. Second, we were interested in whether we could detect differences in DNAm between the smoker and nonsmoker using DNAm level derived from Nanopore sequencing at sites within the targeted regions. We applied Fisher’s test to compare the proportion of methylated reads between the two samples at 514 sites profiled at sufficient read depth (> 10) in both samples (**Supplementary Table 3**). Twelve CpGs had a Bonferonni adjusted significant p-value, 10 in the AHRR region and 2 in the chromosome 2 intergenic region. All 12 CpGs were hypomethylated in the smoker, with a mean difference of −0.53. The power to detect effects in the sequencing based DNAm analyses depends not only on the magnitude of effect but also the read depth at that position(3). We wanted to determine, whether we had potentially missed associations due to limited sequencing coverage. Comparing the level of significance against total read depth across both samples, we observed that the lowest combined read depth of a significant site was 44, more than double our read depth filter of at least 10 in both samples, indicating that at some sites we were not sufficiently powered (**Supplementary Figure 6**). Next, we compared our results with an EWAS of tobacco smoking based on participants from the UK Household Longitudinal Study (UKHLS) who had whole blood DNAm profiled using the EPIC array, to confirm whether we could validate and refine previously reported associations (**Supplementary Table 4**). There were 39 CpGs tested with both platforms and only one site was significant in both analyses, (**Figure 3A**). However, in general, for sites significant in the EPIC EWAS, the nanopore sequencing data demonstrated the same direction of effect as that reported in the EPIC array EWAS even if it was not significant (**Figure 3B**). To establish whether the lack of overlap of significant associations was due to insufficient read depth in the nanopore sequencing data, we compared the EPIC array p-values with read depth and indeed, of the significant sites from the EPIC EWAS, the one that was also significant in our nanopore analysis had the highest read depth (**Figure 3C**). Therefore, we conclude that our inability to rediscover all previously reported smoking sites is due to limited power despite enrichment in our targeted regions.

**Figure 2.**
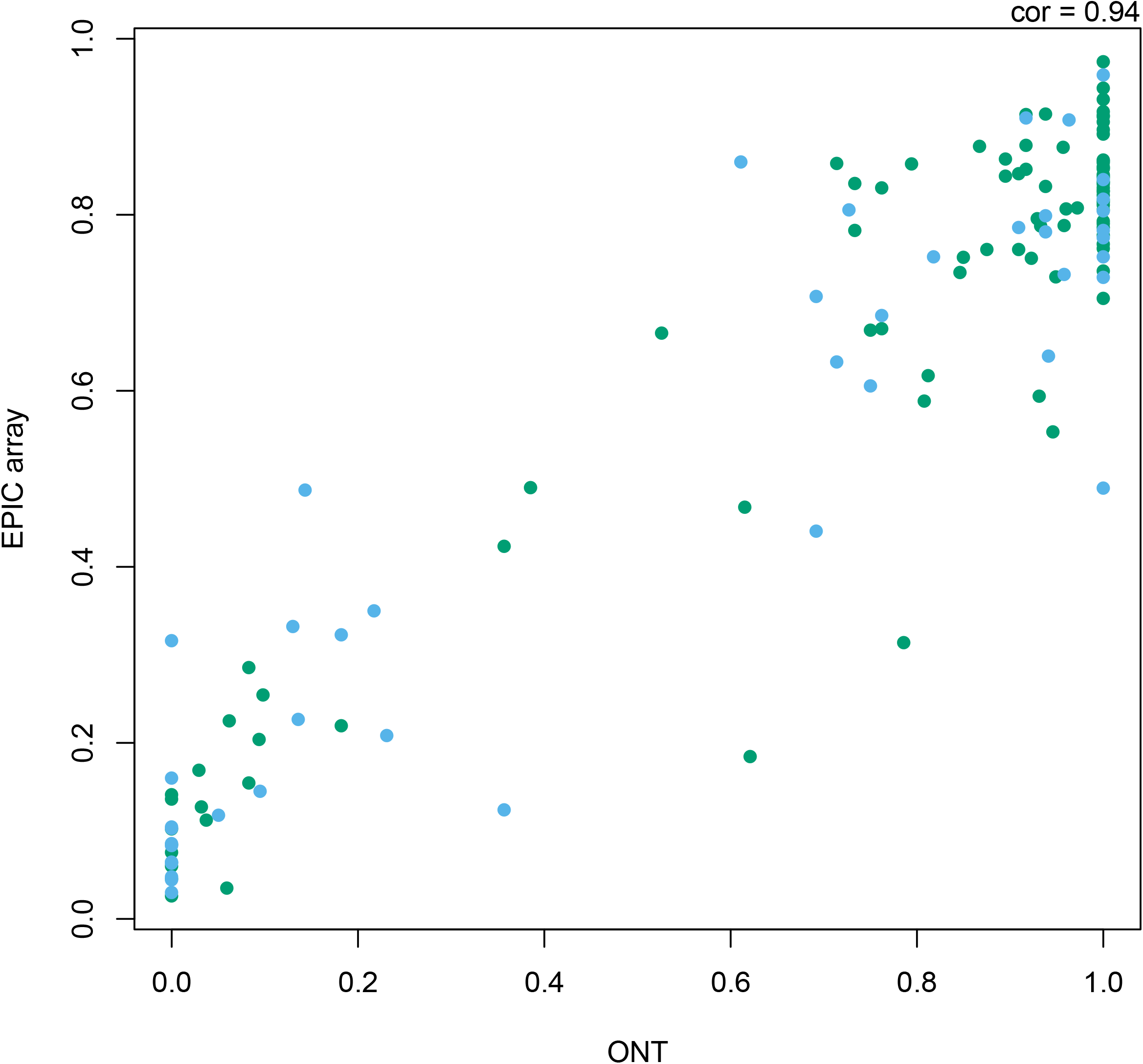
Scatterplot of DNAm quantified using ONT sequencing and Illumina EPIC arrays. Plotted is the level of DNAm estimated from Nanopore reads using Nanopolish (x-axis) and Illumina EPIC arrays (y-axis) for all sites within the three targeted regions which were profiled using both platforms combined across both samples. The colour of the point differentiates the two samples (i.e. smoker and non-smoker).

**Figure 3.**
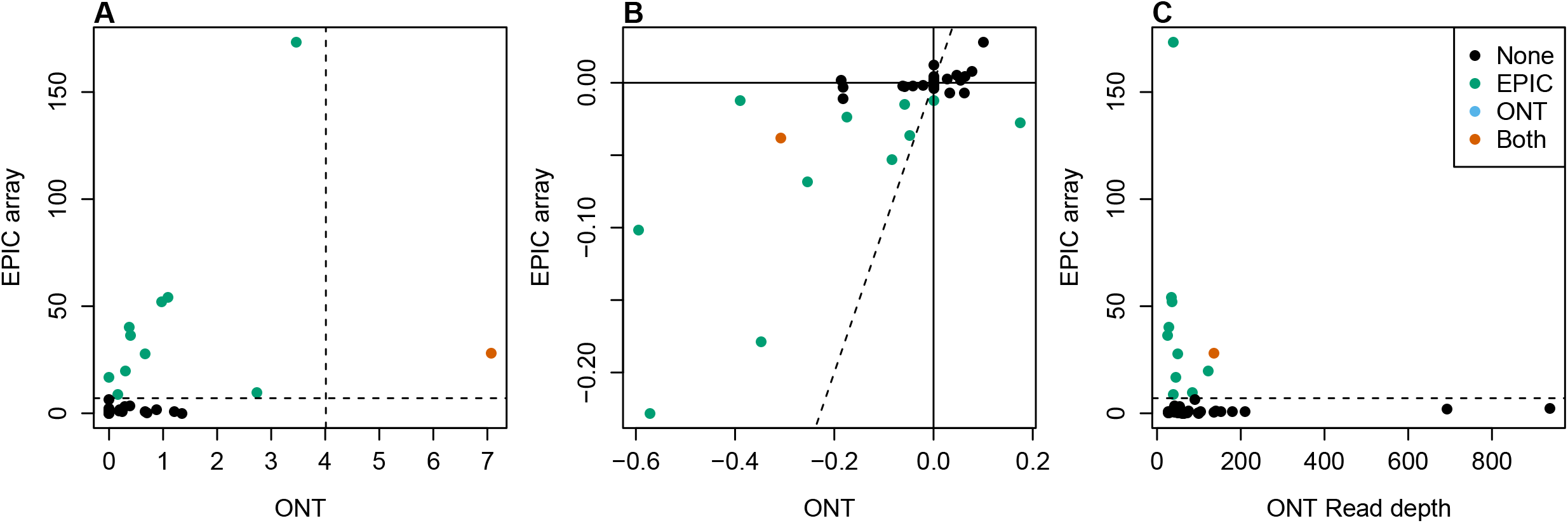
Comparison of nanopore sequencing and Illumina EPIC array to identify differential methylation sites associated with smoking. Across the 34 CpGs tested with both Nanopore sequencing and EPIC array for associations with current smoking status, A) scatterplot of −log10 p-values from EWAS comparing smoker(s) and non-smoker(s) using Nanopore sequencing (x-axis) and EPIC array (y-axis). B) Scatterplot of the (mean) difference in DNAm between smoker(s) and non-smoker(s) estimated using Nanopore sequencing (x-axis) or EPIC array (y-axis). C) Scatterplot of total read depth in Nanopore sequencing (x-axis) and −log10 p-values from EWAS comparing smokers and non-smokers using EPIC array (y-axis). In all panels the colour of the point indicates with which technology a significant difference was detected.

Eleven of the smoking associated significantly CpGs we detected with Nanopore sequencing are not present on the EPIC array and therefore represent novel associations. Looking at the genomic position of these, all ten of the significant sites located within AHRR are intronic, with nine annotated to the same intron (**Supplementary Figure 7**). Furthermore, we observed that 6 of these CpGs clustered within 400bp (**Figure 4**) and overlap with cg05575921, typically the site on the EPIC array with the most replicated association due to its large magnitude of effect (7, 9, 11, 13, 16, 17). For further functional annotation, we downloaded the predicted regulatory functions from ChromHMM(18) for blood, and found that these six CpGs were located in a bivalent enhancer region, while the other CpGs in the AHRR region were located in repressed regions. The two significant CpGs located on chromosome 2 are *1kb apart within the same CpG island and lie within a broader region of associated sites identified with the EPIC array (**Supplementary Figure 8**).

**Figure 4.**
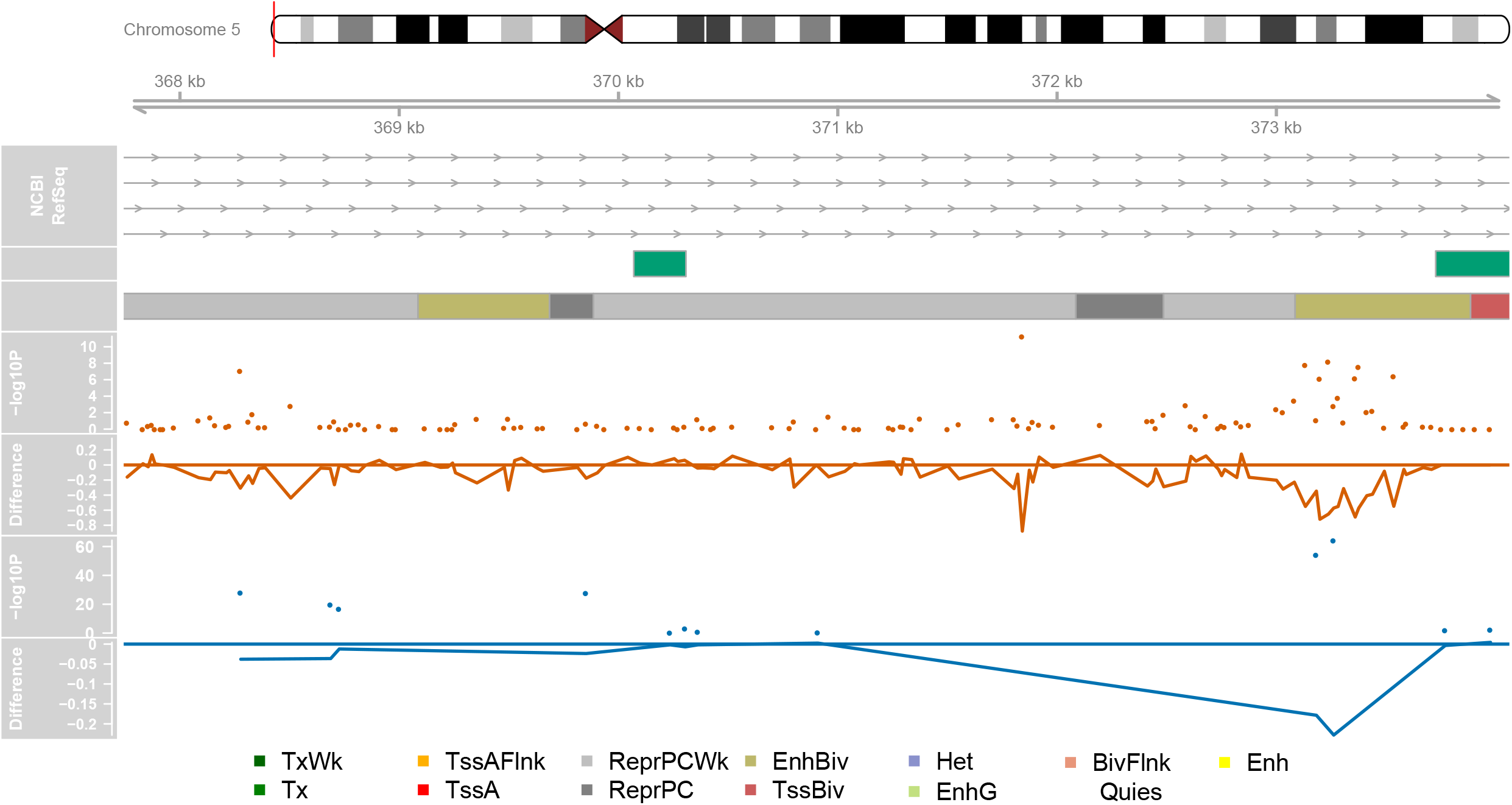
Genomic distribution of AHRR CpGs with significantly different proportions of DNAm associated with smoking. Depicted is part of the targeted genomic region on chromosome 5 containing AHRR where a cluster of significant CpGs were identified. Shown from top to bottom is the gene locations (exons and introns) for different transcripts, CpG islands locations (green boxes), chromHMM predicted chromatin annotations from the 15 state model for blood (E062) where the colour of the box indicates the type of regulatory region as conferred in the legend at the bottom of the panel, a (orange) Manhattan plot of the −log10 P-values from the Fisher’s test of the Nanopore sequencing estimated DNAm proportions comparing a smoker and non-smoker, a (orange) line graph of the estimated difference in DNAm proportion between the smoker and non-smoker from the Nanopore data, a (blue) Manhattan plot of the −log10 P-values from the EPIC array EWAS of current smoking status and a (blue) line graph of the estimated mean difference in DNAm proportion between smokers and non-smokers estimated with the EPIC array.

An additional benefit of profiling DNA methylation with long read sequencing is the ability to determine whether correlated methylation status between neighbouring sites occurs non-randomly. We calculated an adapted version of the linkage disequilibrium statistic D’ between pairs of sites profiled in the same read to quantify whether the co-occurrence of methylation status was greater than expected by chance, given the proportion of methylation at those sites. While there were pairs of CpGs that had the same methylation status within a read, this is not as extensive or prevalent as is typically observed across genetic variants. First, there was no evidence of decay in these relationships as the distance between sites increased (**Supplementary Figure 9)**. Second, the sites did not segregate cleanly into “blocks” of highly correlated methylation calls (**Supplementary Figure 10**). Instead, it was seemingly random pairs of sites that were highly co-ordinated. Considering just the subset of 12 sites with significant differences associated with smoking status, we did not see any evidence for the methylation status of these to co-occur with the same read in a non-random manner (**Supplementary Figure 11**).

## Discussion

In this study, we performed the first quantitative assessment of Nanopore sequencing for epigenetic epidemiology by deriving DNAm profiles from native DNA and comparing with profiles generated with the current standard microarray technology (EPIC array). Our analyses focused on three genomic regions, selected from previous EWAS of tobacco smoking(7, 9, 13, 16), which we targeted using CRISPR gRNAs, to test whether the sensitivity of DNAm quantification from Nanopore data is sufficient to rediscover these associations. The correlation between technologies was very high and the estimated level of DNAm accurate enough to detect significant differences between a heavy smoker and non-smoker at genomic loci reported in previous analyses with much larger sample sizes. One of the key advantages of using sequencing to profile DNAm is the greater spatial resolution of signals across the genome. For example, in our data, we had *9 fold more sites across the AHRR gene compared to the content of the EPIC array, enabling us to discover additional novel loci in this region associated with smoking that have not previously been analysed. This has the potential to advance our understanding of the role of aberrant differential methylation in the aetiology of complex diseases by providing complete coverage of the region rather than being limited to a predefined subset of sites that may or may not capture the complete extent of methylomic variation in that region. A specific utility of long read sequencing over both microarrays and short read sequencing is the ability to characterise whether methylation status is coordinated across CpGs from the same genomic region by quantifying the proportion of reads with concordant methylation calls was greater than expected by chance. We found that high correlations between neighbouring sites were the exception, meaning that existing studies likely do not capture much information about unmeasured sites and that it is unlikely that the imputation of DNA methylation levels will be as effective as it is for studies of genetic variation. Altogether, this reiterates the need to empirically profile DNA methylation using technologies that are genuinely genome-wide. Improving the spatial resolution of DNAm quantification will clarify the genomic region over which differential methylation occurs, permitting better functional annotation and enabling biological inferences.

While our data show great promise for the role of Nanopore sequencing in studies of DNAm, it also highlighted some issues that will affect how it should be used. DNAm is quantified as a proportion and when using sequencing reads it is calculated as the fraction of methylated reads to total number of reads at that position. The accuracy of the quantification is therefore, dependent on the sequencing depth at that position (i.e. the denominator in the calculation). However, as in a typical sequencing experiment the majority of DNAm sites are captured by a handful of reads, while the total number of CpGs covered can be many orders of magnitude higher than a study based on microarrays, only a minority are profiled adequately for any downstream statistical analysis. To improve the likelihood of detecting differences, we used a targeted approach based on CRISPR/Cas9 methodology(6) to enrich for CpGs in three specific genomic regions of varying size. This limited our ability to detect novel DMPs to those located with the regions that are already implicated. After filtering DNAm sites for minimum coverage of 10 reads, only sites within our targeted regions where retained. Despite the high proportion of off-target reads, the mean read depth across the genome was insufficient for accurate quantification, highlighting the necessity of an enrichment step.

The methodology we present is applicable to any genomic region, and we have shown that it is feasible to consider multiple targets in a single experiment. It is hard to predict from our data whether the magnitude of coverage enrichment we report would be replicated if we had included more targeted regions, or whether we would have seen higher levels of enrichment if we had considered fewer regions. For regions that are smaller than the typical read length only gRNAs at the start and end are needed, whereas for target regions larger than the typical read length (e.g. the AHRR region) additional gRNAs tiled across the region are required. The way the gRNAs combine is random If there are lots of gRNAs too close together multiple small fragments may be produced. If they are too far apart, and the reads do not span the full extent of the gap between then, then an important part of the region may not be adequately covered. Across the AHRR region, where the gRNAs were located close together, we saw greater levels of enrichment in terms of number of reads. All the gRNAs we included produced fragments but the performance was variable, with the enrichment around GFI1 less successful in part because one of the gRNAs was incorrectly orientated. Therefore, we would recommend doubling up on gRNAs to protect against the variable efficiency and to ensure adequate coverage at particularly important regions. To maximise the probability of the “correct” gRNAs pairing up on the same fragment we ran the targeted regions in different pools, with multiple pools for the AHRR region consisting of different gRNAs. It should be noted that one technical limitation of this method for profiling DNA methylation is that it requires ten times the DNA input per pool of gRNAs, compared to microarrays, and therefore increasing the number of regions and therefore pools will have an effect on the quantity of DNA required for sequencing.

As well as targeting specific regions of the genome where we knew differences existed, we additionally chose a phenotype associated with large effects, such that differences should be detectable even if the sensitivity is low. While this strategy was effective, it is unclear from our analysis how viable Nanopore sequencing will be for detecting smaller differences between groups. Even within our targeted regions, there were a number of previously reported sites where we did not detect statistically significant differences, which we hypothesise is due to insufficient read depth despite target enrichment. Despite the experiment successfully enriching the data for coverage with our three targeted regions, it should be noted that the vast majority of reads were located outside of these and randomly distributed across the genome, meaning they were excluded from the analysis due to low coverage. Improving the efficiency of the enrichment will be the key to establishing this approach for studying a broad range of complex diseases and phenotypes.

Another important factor for study design is sample size. EWAS based on microarrays require large sample sizes to robustly detect differences after adjusting for the penalty of testing sites across the genome(15, 19), with the exact size of the sample dependent on the magnitude of the effect associated with the phenotype under investigation(20). When quantifying DNAm through sequencing based approaches, statistical power additionally related to sequencing depth, which can be increased by either profiling more samples, or sequencing the samples you have more deeply. Both approaches have financial and practical implications. For Nanopore sequencing, currently there is no methodology to enable multiple samples to be profiled on a single flowcell, limiting the total sample size. In this study we only profiled two individuals, one heavy smoker and one non-smoker, while the quantity of sequencing data made up for the lack of samples, this may have affected which differences we were able to detect if there is any inter-individual variation in terms of which sites are affected or the magnitude of the difference. In order to capture the sites, which we did not rediscover, either additional samples or more flowcells would likely be required.

As well as the experimental methodology, the data analysis pipeline requires careful thought. Calling DNAm from Nanopore sequencing data is a computational challenge to classify methylated and unmethylated cytosines based on the electrical signal emitted as the DNA passes through the pore with a number of algorithms developed for this purpose. In this study we implemented just one of these algorithms, Nanopolish(4), which was found to be consistently one of the most accurate and concordant with the other best performing methods, across a range of genomic contexts, as well as the least computationally intensive(21). One limitation of this algorithm is that it only considers CpGs and ignores DNAm at cytosines in other genomic contexts. This is of little consequence for our comparison with the EPIC array, as it also predominantly focuses on CpG sites, but means that there is another layer of resolution in the DNAm profiles we have not considered. As methods for calling additional DNA modifications are validated, an additional advantage will be the ability to call multiple epigenetic marks from a single sequencing run(22). This would be especially beneficial for studies of the brain where DNA hydroxymethylation is abundant(23).

In summary, our data indicates that Nanopore sequencing is a valid option for identifying multiple CpGs across the genome that are associated with large differences in DNAm between groups. It has the potential to fine map associations detected with existing microarray platforms by validating previous associations and identifying novel loci and in this way advance our understanding of the role differential methylation plays in the aetiology of complex disease.

## Materials and Methods

### Samples

Matched (age/sex) samples, including one heavy smoker and one non-smoker, were obtained from the Exeter 10,000 and Peninsula Research Bank (EXTEND/PRB), an ethically approved biobank providing access to anonymised DNA/RNA/Urine/Plasma/Serum. Samples are stored at −80°C. (https://exetercrfnihr.org/about/exeter-10000-prb/). The EXTEND/PRB is housed and managed within the NIHR Exeter Clinical Research Facility (Exeter CRF).

### Design of CRISPR/Cas9 gRNAs

Three genomic regions that contained robust smoking-associated differentially methylated sites were selected as the target regions (**Table 1**). For each region, we designed two gRNAs for the start and two gRNAs for the end of the region. For the AHRR region which is 140kb and longer than the average read generated by Nanopore sequencing, six additional guides were designed, tiled across the region. We used the Alt-R^®^ CRISPR-Cas9 system from Integrated DNA Technologies (IDT). The gRNACRISPR were designed using software available through the IDT website, selecting those with the optimal predicted efficiency and specificity scores. In total 18 gRNAs were included, the details of which are in **Supplementary Table 1**. The gRNA were ordered as CRISPR RNA (crRNA), to allow the formation of RNP with IDT tracRNA and Cas9 protein.

### Cas9 cleavage and library prep

The CRISPR Cas9-mediated target enrichment was carried out in line with the ONT protocol (Cas-mediated PCR-free enrichment, ENR_9084_v109_revM_04Dec2018). In accordance with the tiling approach described by ONT, the crRNAs were diluted to 100μM in TE pH 7.5 (IDT). crRNA were split across five pools at equimolar concentrations, with two pools for the larger AHRR region, one pool each for the two small regions, and a final pool with all 18, as detailed in **Supplementary Table 1**.

1μl of each crRNA pool (100μM) was combined with 1μl of tracrRNA (100μM) and 8μl IDT duplex buffer. This solution was incubated at 95°C for 5 minutes in a Veriti™ 96-well thermal cycler (Applied Biosystems™) which then ramped down slowly to 25°C. To form functional CRISPR ribonucleoproteins (RNPs), 3μl of this annealed crRNA/tracrRNA was incubated with 0.3μl 62μM HiFi Cas-9 (IDT), 3μl 10X NEB CutSmart^®^ Buffer (New England BioLabs) and 23.7μl nuclease free water for 30 minutes at room temperature.

Prior to Cas9 cleavage, human genomic DNA (gDNA) was dephosphorylated using Calf Intestinal Phosphatase. Briefly, 25μg of gDNA (5μg per crRNA pool) was diluted in 15.6μl 10X CutSmart^®^ Buffer (New England BioLabs) and 15.6 μl Calf Intestinal Phosphatase (5U/μl) (New England BioLabs) and was incubated for 10 minutes at 37°C, 2 minutes at 80°C and then held at 20°C in a Veriti™ 96-well thermal cycler (Applied Biosystems™) until cleavage.

For cleavage 10μl of each CRISPR RNP pool was combined with 5μg dephosphorylated gDNA, 1μl 10mM dATP (New England BioLabs) and 1μl NEB Taq Polymerase (New England BioLabs) in order to achieve targeted gDNA cleavage and dA tailing of cleaved products. The reaction was incubated at 37°C for 30 minutes, 72°C for 5 minutes and then held at 4°C in a Veriti™ 96-well thermal cycler (Applied Biosystems™). Adaptors from the LSK-109 sequencing kit (Nanopore) were then ligated onto 42ul of pooled CRISPR cleaved DNA using NEBNext Quick T4 Ligase (New England BioLabs) and incubated at room temperature for 20 minutes. 1 volume of TE was then added and 0.3X AMPure XP bead purification (Beckman), using 2 x 250ul SFB buffer washes in place of Ethanol. The library was eluted in 13ul of EB buffer (Nanopore) and loaded on a Nanopore Mk1b sequencer with MinION r9.4.1 flowcell and run for 24h.

### DNA methylation EPIC array

DNAm data for the two samples included in this study was generated as part of a larger project profiling > 1,200 individuals from the EXTEND cohort. The EZ-96 DNA Methylation-Gold kit (Zymo Research; Cat No# D5007) was used for treating 500 ng of DNA from each sample with sodium bisulfite. DNA methylation data were generated using the Illumina Infinium HumanMethylationEPIC BeadChip (“EPIC array”) array. Raw data was processed using the wateRmelon package (24) and put through a stringent quality control pipeline that included the following steps: (1) checking methylated and unmethylated signal intensities and excluding poorly performing samples; (2) assessing the chemistry of the experiment by calculating a bisulphite conversion statistic for each sample, excluding samples with a conversion rate <80%; (3) identifying the fully methylated control sample was in the correct location; (4) multidimensional scaling of sites on the X and Y chromosomes separately to confirm reported sex; (5) using the 59 SNPs on the Illumina EPIC array to check for sample duplications; (6) use of the pfilter() function in wateRmelon to exclude samples with >1 % of probes with a detection P value □>□ 0.05 and probes with >1 % of samples with detection P valued□>□0.05; (7) normalisation of the DNA methylation data was performed using the dasen() function in wateRmelon(24); (8) samples that were dramatically altered as a result of normalization were excluded on the basis of the difference between the normalized and raw data; and (9) removal of cross-hybridising and SNP probes (2, 25).

### Statistical analysis

Base calling was performed using GUPPY (Version 4.0.11, high accuracy model) to generate FASTQ sequencing reads from the electrical data. Reads were aligned to the human reference genome (hg38) using Minimap2(26). Aligned reads were then filtered to primary alignments and reads of high quality using samtools. Nanopolish(5) was then used to call DNAm from individual reads which were then aggregated into estimatess of the level of DNAm by counting the proportion of methylated reads to the total number of reads at position using the script provided in Nanopolish. To compare the level of DNAm at individual sites between the smoker and non-smoker a Fisher’s test was used to compare the proportion of methylated reads between the two samples. Significant sites were identified after adjusting the p-values for the total number of sites tested (514) using the Bonferroni method. Genomic region plots were generated using the Gviz package(27). To profile whether concordant methylation status at neighbouring sites occurs non-randomly, we adapted the linkage disequilibrium statistic D’ to quantify whether the co-occurrence of methylation at pairs of sites within a read was greater than expected by chance, given the proportion of methylation at those sites. For pairs of sites that were profiled in same read, in at least 10 reads across both samples, D was calculated as the proportion of reads where the methylation status (either methylated or unmethylated) was consistent at both sites minus the probability of the status being consistent given the proportion of methylation at each site (see equation below). D was then standardized to D’, by dividing it by its theoretical maximum.

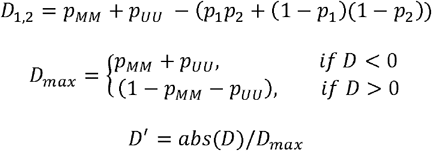

Where *p_MM_* is the proportion of reads where both sites are methylated, *p_UU_* is the proportion of reads where both sites are unmethylated, *p*_1_, *p*_2_ are the proportion of reads that are methylated at sites 1 and 2 respectively. Heatmaps of ‘linkage’ statistics between pairs of DNAm sites were generated using the LDheatmap package(28).

All analysis was performed with the R statistical language version 3.6.3. All analysis scripts are available at https://github.com/ejh243/ONTMethCalling.

### EPIC array based EWAS of tobacco smoking

The British Household Panel Survey (BHPS) began in 1991, and in 2010 was incorporated into the larger UK Household Longitudinal Study(29) (UKHLS; also known as Understanding Society) which is a longitudinal panel survey of 40,000 UK households from England, Scotland, Wales and Northern Ireland. DNAm was profiled in DNA extracted from whole blood for 1,170 individuals who were eligible for and consented to both blood sampling and genetic analysis, had been present at all annual interviews between 1999 and 2011, and whose time between blood sample collection and processing did not exceed 3 days. Eligibility requirements for genetic analyses meant that the epigenetic sample was restricted to participants of white ethnicity. 500ng of DNA from each sample was treated with sodium bisulfite, using the EZ-96 DNA methylation-Gold kit (Zymo Research, CA, USA). DNAm was quantified using the Illumina Infinium HumanMethylationEPIC BeadChip (Illumina Inc, CA, USA) run on an Illumina iScan System (Illumina, CA, USA) using the manufacturers’ standard protocol. Samples were randomly assigned to chips and plates to minimise batch effects. Quality control, pre-processing and data normalisation were carried out using the bigmelon package(30) following a standard pipeline(31).

Smoking status was derived from interview data and the response to the question “Do you smoke cigarettes now?” to classify as either a current or non-smoker. In total 1,113 participants were included in the EWAS of current smoking status. To identify sites where DNAm was significantly different between smokers and non-smokers, a linear model was fitted using the limma R package(32) for all sites on the EPIC array controlling for age, sex, six cell type proportions (CD8T, CD4T, NK, Bcell, Mono, Gran) (33, 34) and plate as a potential source of technical variation.

## Acknowledgements

This work has funded by a Brain and Behaviour Foundation Young Investigator Award to EH [26288] and utilised equipment funded by the UK Medical Research Council (MRC) Clinical Research Infrastructure Initiative (award number MR/M008924/1). EH, LS and JM are supported by MRC grant K013807 to JM. The authors would like to acknowledge the use of the University of Exeter High-Performance Computing (HPC) facility. The NIHR Exeter Clinical Research Facility is a partnership between the University of Exeter Medical School College of Medicine and Health, and Royal Devon and Exeter NHS Foundation Trust. This project is supported by the National Institute for Health Research (NIHR) Exeter Clinical Research Facility.□ The views expressed are those of the author(s) and not necessarily those of the NIHR or the Department of Health and Social Care.

## Abbreviations

crRNA: CRISPR-Cas9 CRISPR RNAs
DNAm: DNA methylation
DMP: differentially methylated region
gDNA: genomic DNA
IDT: Integrated DNA Technologies
ONT: Oxford Nanopore Technologies
RNPs: ribonucleoproteins

