## Supplementary Figure for "Evaluation of nanopore sequencing for epigenetic epidemiology: a comparison with DNA methylation microarrays"

**Supplementary Figure 1.** Distribution of nanopore sequence reads across GFI1 region. Depicted is the targeted genomic region on chromosome 1 containing GFI1 and the location of the guideRNAs and sequencing reads. Shown from top to bottom is the gene locations (exons and introns) for different transcripts, CpG islands locations (green boxes), target position of the guideRNAs where the grey arrow indicates the strand targeted, a histogram of the total number of nanopore reads overlapping each position , and the location of the actual reads at the bottom. Note that due to the number of reads, only a subset are included to give a representative view of read mappings.

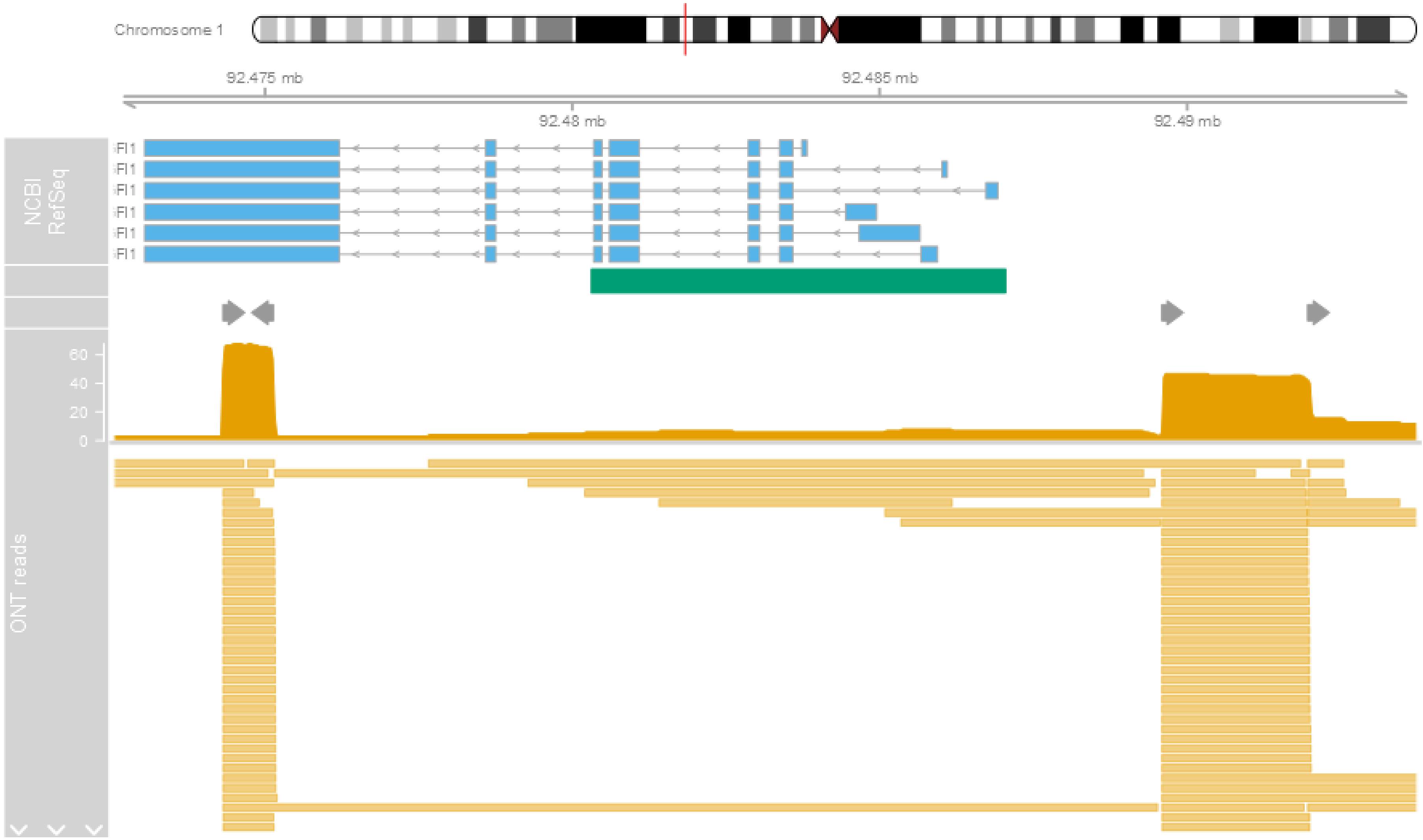

**Supplementary Figure 2.** Distribution of nanopore sequence reads across chromosome 2 region. Depicted is the targeted genomic region on chromosome 2 and the location of the guideRNAs and sequencing reads. Shown from top to bottom is the gene locations (exons and introns) for different transcripts, CpG islands locations (green boxes), target position of the guideRNAs where the grey arrow indicates the strand targeted, a histogram of the total number of nanopore reads overlapping each position , and the location of the actual reads at the bottom. Note that due to the number of reads, only a subset are included to give a representative view of read mappings.

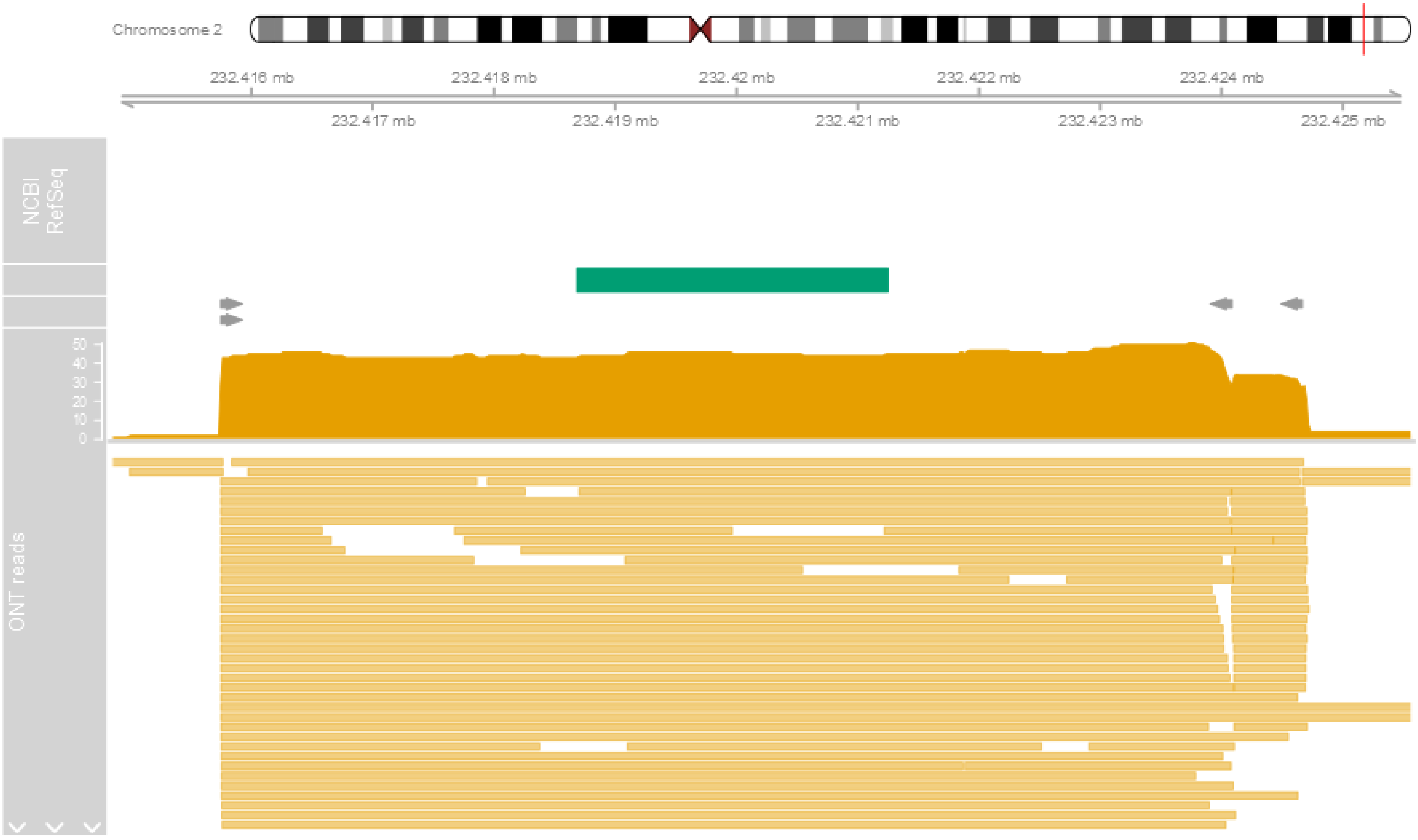

**Supplementary Figure 3.** Barplot showing the distribution of number of reads per chromosome.

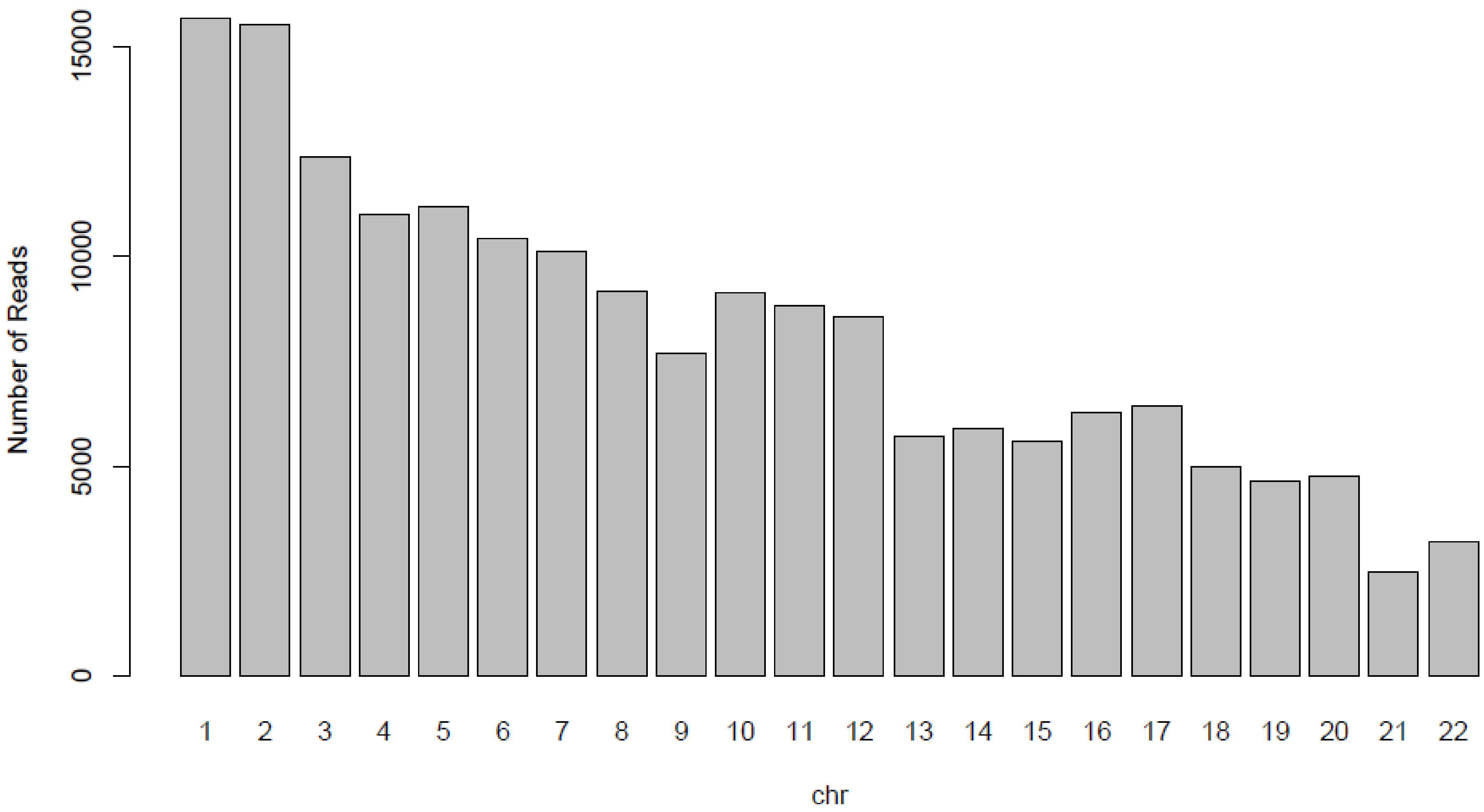

**Supplementary Figure 4.** Histograms showing the distribution of proportion of each of the three-targeted regions covered by each read.

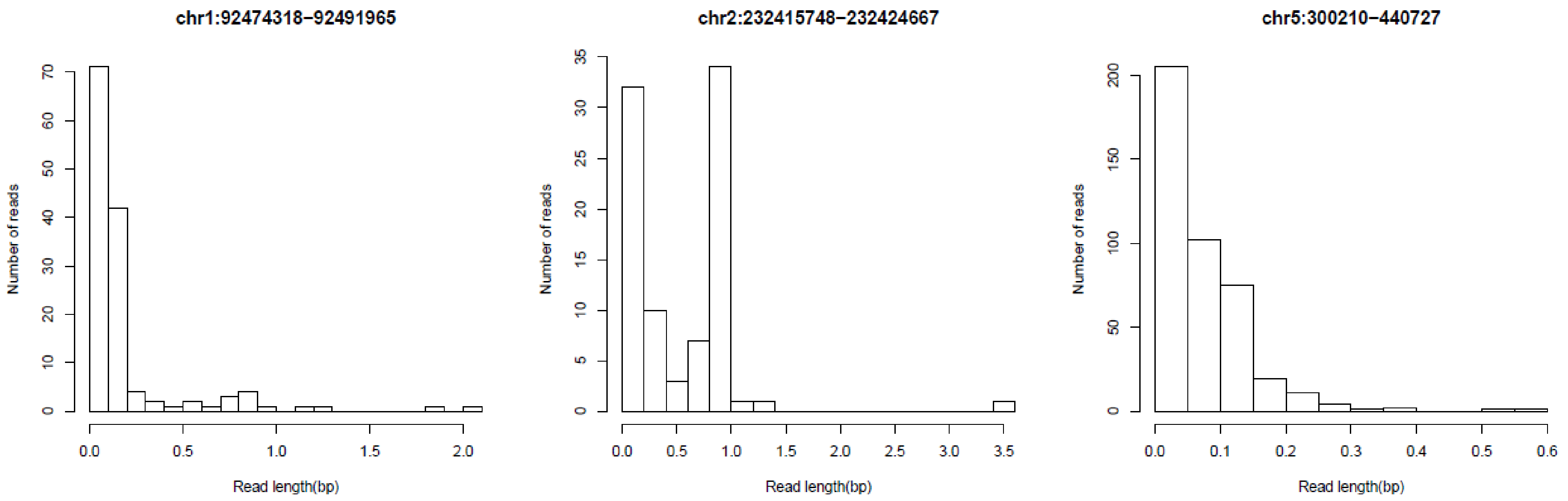

**Supplementary Figure 5.** Histograms showing the distribution of read lengths within each of the three-targeted regions.

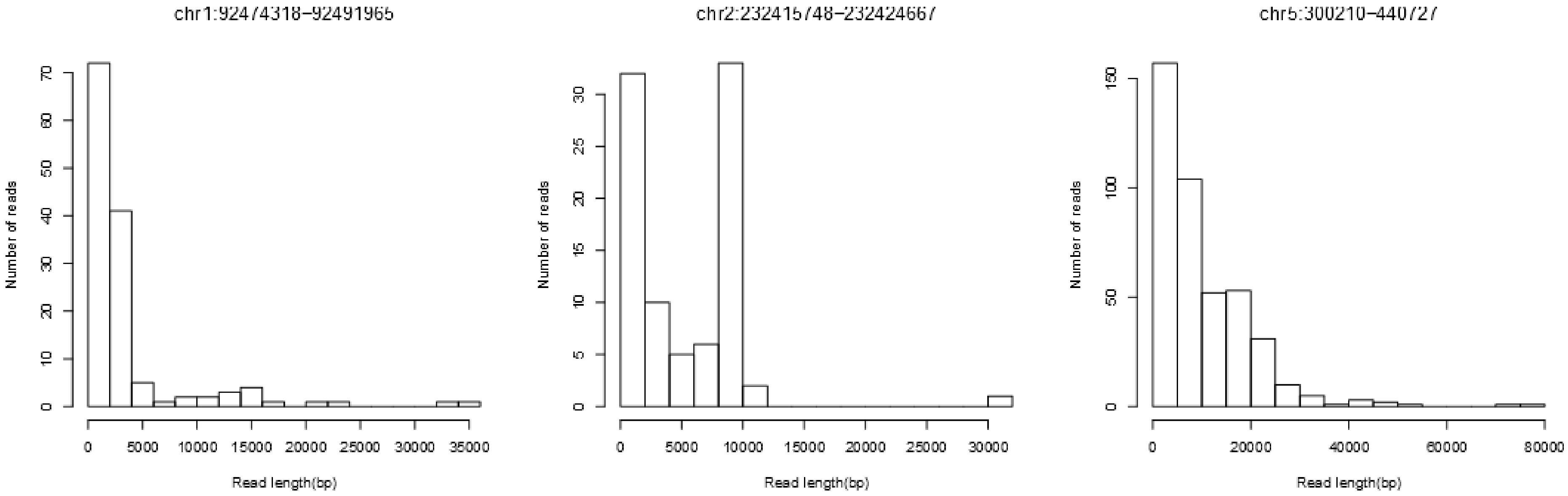

**Supplementary Figure 6.** Scatterplot of –log10 P-value from Fisher's test comparing proportion of methylated reads between the smoker and non-smoker against combined read depth across both samples. The panel on the right is a zoomed in version of the panel on the left. The horizontal dashed line indicates the multiple testing adjusted significance threshold.

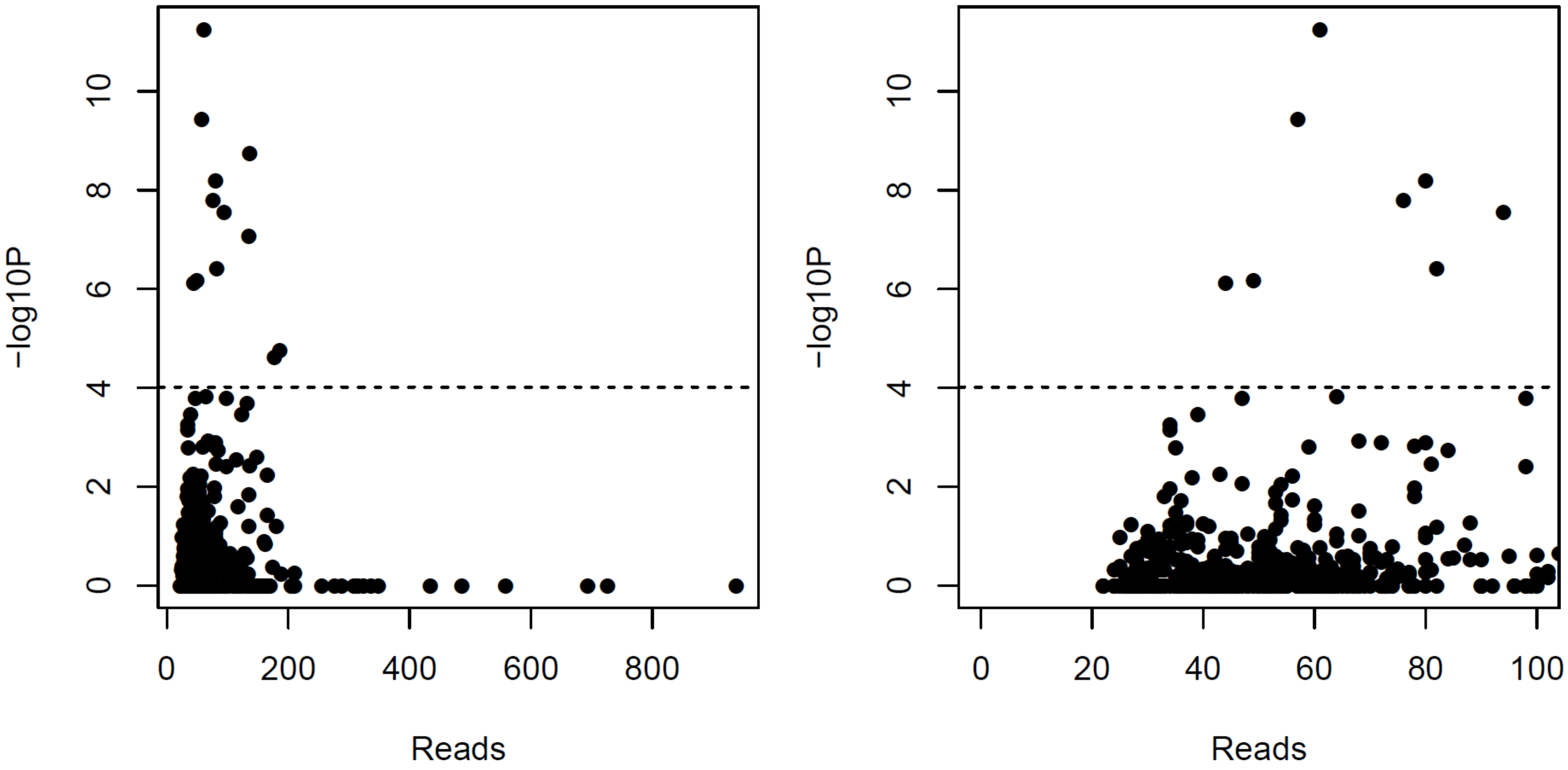

**Supplementary Figure 7.** Genomic distribution of AHRR CpGs with significantly different proportions of DNAm associated with smoking. Depicted is the targeted genomic region on chromosome 5 containing AHRR. Shown from top to bottom is the gene locations (exons and introns) for different transcripts, CpG islands locations (green boxes), chromHMM predicted chromatin annotations for E062 where the colour of the box indicates the type of regulatory region as conferred in the legend at the bottom of the panel, a (orange) Manhattan plot of the –log10 P-values from the Fisher's test of the nanopore estimated DNAm proportions comparing a smoker and non-smoker, a (orange) line graph of the estimated difference in DNAm proportion between the smoker and non-smoker from the nanopore data, a (blue) Manhattan plot of the –log10 P-values from the EWAS array EWAs of current smoking status and a (blue) line graph of the estimated mean difference in DNAm proportion between smokers and non-smokers estimated with the EPIC array.

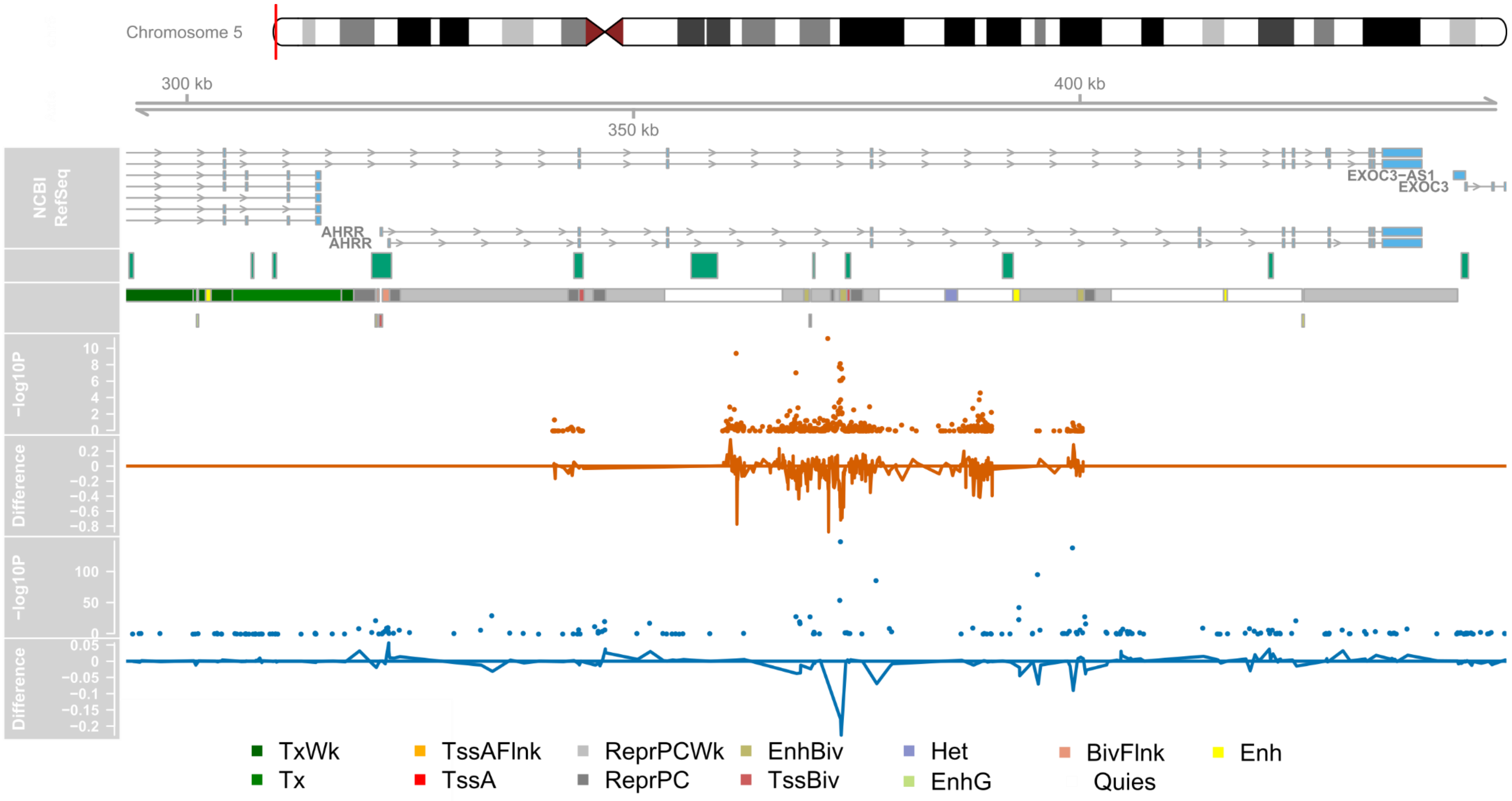

**Supplementary Figure 8.** Genomic distribution of chromosome 2 CpGs with significantly different proportions of DNAm associated with smoking. Depicted is the targeted intergenic genomic region on chromosome 2. Shown from top to bottom is the gene locations (exons and introns) for different transcripts, CpG islands locations (green boxes), chromHMM predicted chromatin annotations for E062 where the colour of the box indicates the type of regulatory region as conferred in the legend at the bottom of the panel, a (orange) Manhattan plot of the –log10 P-values from the Fisher's test of the nanopore estimated DNAm proportions comparing a smoker and non-smoker, a (orange) line graph of the estimated difference in DNAm proportion between the smoker and non-smoker from the nanopore data, a (blue) Manhattan plot of the –log10 P-values from the EWAS array EWAs of current smoking status and a (blue) line graph of the estimated mean difference in DNAm proportion between smokers and non-smokers estimated with the EPIC array.

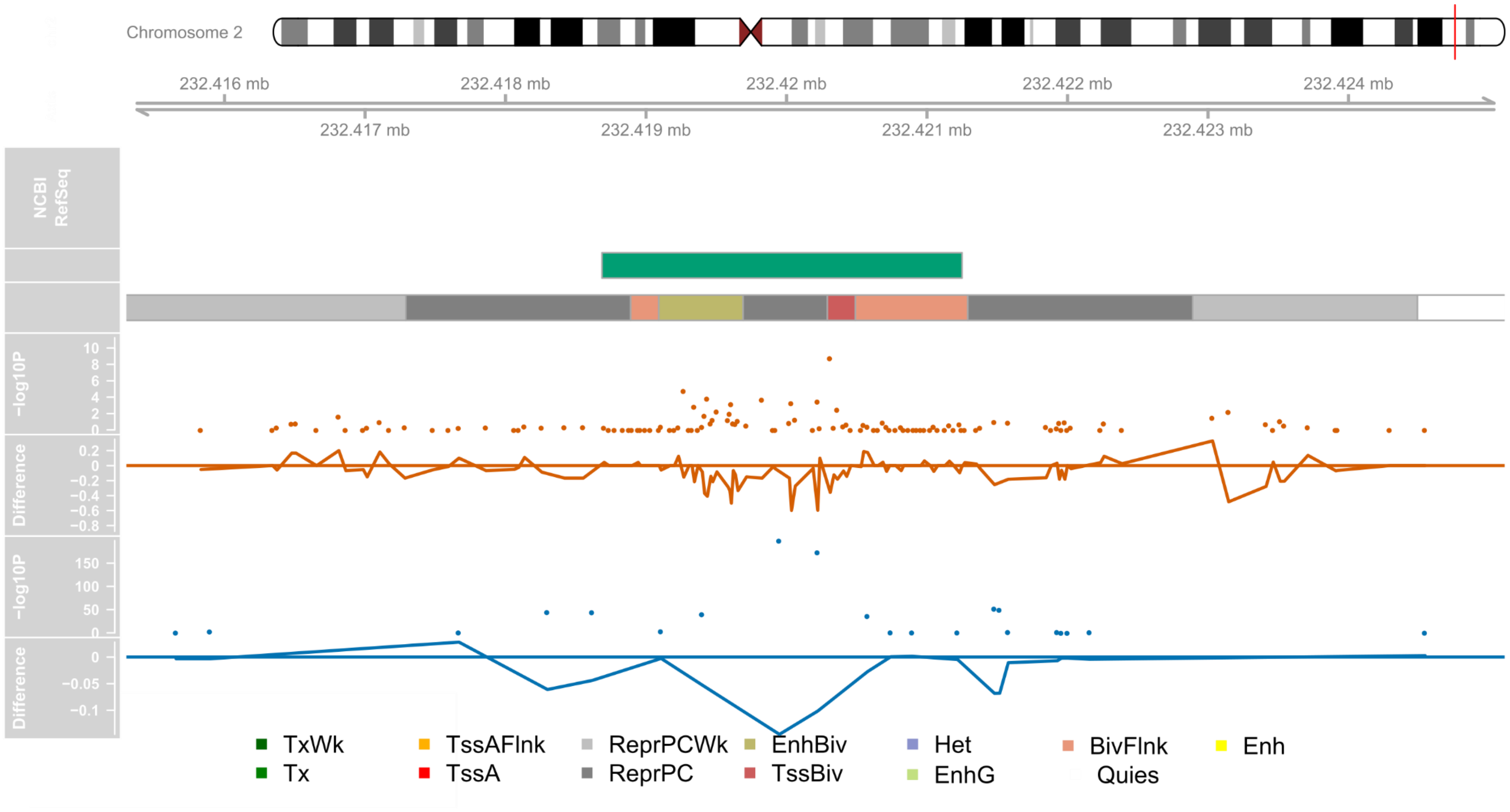

**Supplementary Figure 9.** Scatterplot of DNA methylation status concordance against distance. Points represent all pairs of sites (n = 124,018) within any of the 3 targeted regions with at least 10 reads across both samples. The non-random shared methylation status between pairs of sites profiled within the same read was quantified using an adapted version of the linkage disequilibrium statistics D’ (y-axis). The black represents the moving median for 50 base pair sliding windows.

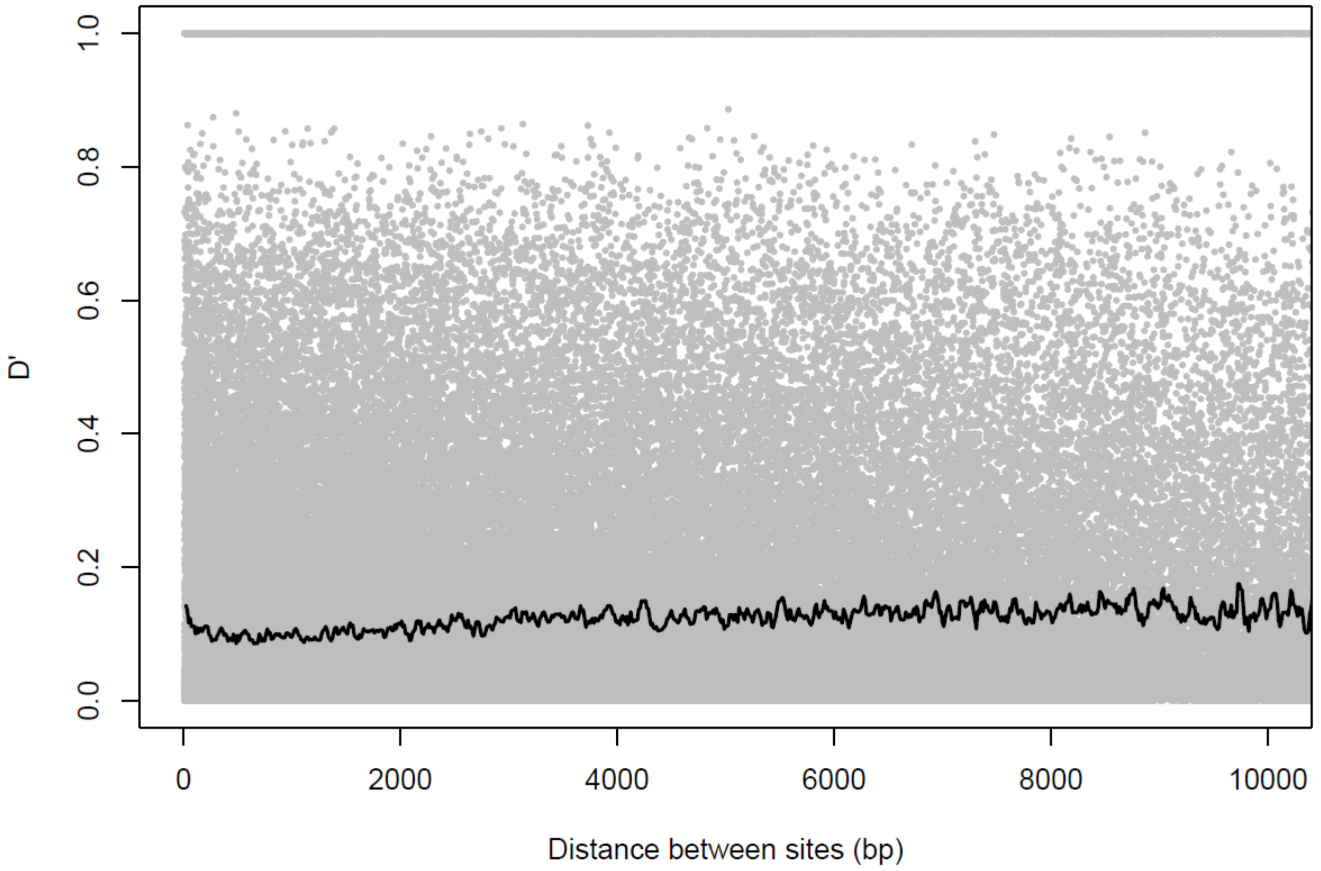

**Supplementary Figure 10.** Heatmap of DNA methylation status concordance between pairs of sites within the targeted AHRR region. Depicted is a subsection of the AHRR targeted region containing 248 DNA methylation sites. The colour of the square represents the D' statistic which quantifies the enrichment of non-randomly co-ordinated methylation status between pairs of sites profiled within the same read.

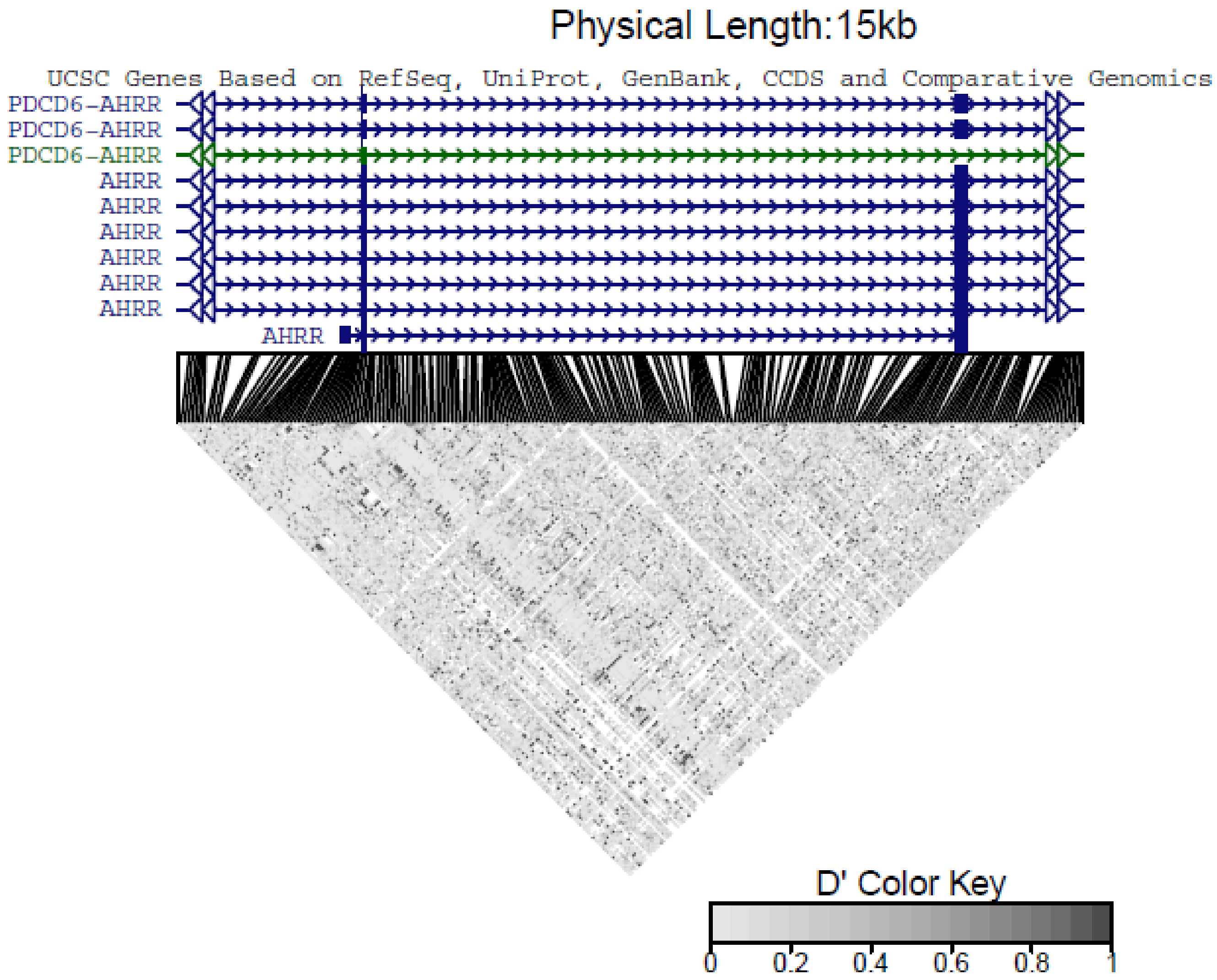

**Supplementary Figure 11.** Heatmap of DNA methylation status concordance between pairs of sites within the regions significantly associated with tobacco smoking. Depicted is a subsection of **A**) chromosome intergenic and **B**) the AHRR targeted region. In the heatmap at the bottom of the plot the colour of the square represents the D' statistic which quantifies the enrichment of non-randomly co-ordinated methylation status between pairs of sites profiled within the same read. Labelled in blue are the DNA methylation sites/regions that were identified as having a significant difference ($P < 9.7x10^{-5}$) between a smoker and non-smoker. At the top of the plot there is a zoomed in Manhattan plot of the –log10 P-values from the Fisher's test of the nanopore estimated DNAm proportions comparing a smoker and non-smoker.

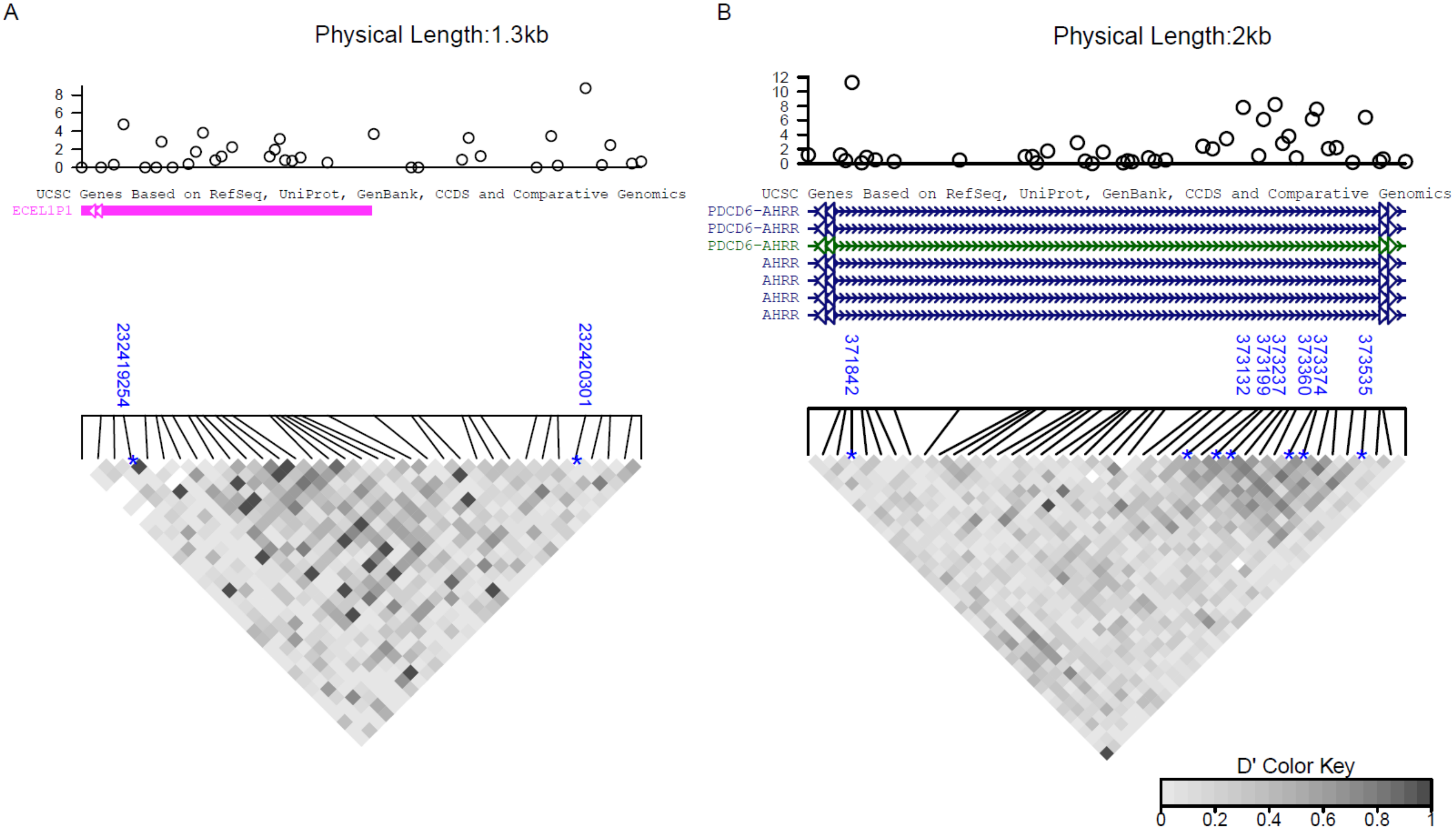
